# MgrA regulates interaction of *Staphylococcus aureus* with mucin

**DOI:** 10.1101/2020.04.09.035261

**Authors:** Connor P. Parker, Nour Akil, Cullen R. Shanrock, Patrick D. Allen, Anna L. Chaly, Heidi A. Crosby, Jakub Kwiecinski, Alexander R. Horswill, Anthony J. Fischer

**Affiliations:** Stead Family Department of Pediatrics, University of Iowa Carver College of Medicine, Iowa City, IA; Department of Immunology & Microbiology, University of Colorado Anschutz Medical Campus, Aurora, CO

**Keywords:** *Staphylococcus aureus*, Mucin, Submucosal Gland, MgrA, Adhesin

## Abstract

**Background:** To defend the lungs, mucus adheres to bacterial cells and facilitates their removal by ciliary transport. Our goals were to measure the affinity of mucus for the respiratory pathogen *Staphylococcus aureus* and identify bacterial genes that regulate this interaction.

**Methods:** *S. aureus* was added to pig tracheas to determine whether it binds mucus or epithelial cells. To quantify its affinity for mucus, we developed a competition assay in microtiter plates. Mucin was added over a dose range as an inhibitor of bacterial attachment. We then examined how transcriptional regulator MgrA and cell wall transpeptidase sortase (SrtA) affect bacterial interaction with mucin.

**Results:** In pig tracheas, *S. aureus* bound mucus strands from submucosal glands more than epithelial cells. In microtiter plate assays, Δ*srtA* failed to attach even in the absence of mucin. Mucin blocked wild type *S. aureus* attachment in a dose-dependent manner. Higher concentrations were needed to inhibit binding of Δ*mgrA*. Co-deletion of *ebh* and *sraP*, which encode surface proteins repressed by MgrA, suppressed the Δ*mgrA* binding phenotype. No differences between Δ*mgrA* and wild type were observed when methylcellulose or heparin sulfate were substituted for mucin, indicating specificity.

**Conclusions:** Mucin decreases attachment of *S. aureus* to plastic, consistent with its physiologic role in host defense. *S. aureus* deficient in MgrA has decreased affinity for mucin. Ebh and SraP, which are normally repressed by MgrA, may function as inhibitors of attachment to mucin. These data show that specific bacterial factors may regulate the interaction of *S. aureus* with mucus.

## Introduction

Mucus is an important defense against respiratory infections. It is a gel-like material comprised of hydrated mucin proteins and inorganic salts produced on epithelial surfaces. In large animals, airway mucus is produced in the form of strands arising from submucosal glands (1-3). Inhaled particles and pathogens can become entrapped in mucus. After the release of mucus strands from submucosal gland ducts or goblet cells, ciliated cells transport mucus together with any particles trapped in the mucus, expelling them from the airway (1). Although mucus is colloquially described as “sticky,” its adhesive properties are seldom quantified. Proper physiologic function of mucus would require that it have some affinity for a diverse repertoire of particles and pathogens, but not excessively adhere to the epithelium or gland duct surfaces. In cystic fibrosis (CF), mucus may interact with pathogens, but it fails detach from gland ducts and is unable to be carried away by ciliary beating. This results in impaired transport of mucus out of the airway (3). In CF pigs, mucus strands fail to detach from airway submucosal glands and becomes pathologically immobile. Thus, it has been hypothesized that mucus stasis may contribute to the establishment of the chronic bacterial infections in CF.

*S. aureus* is a typical pathogen in CF, as approximately 70% of people with CF are infected with the bacteria (4, 5). Among CF pathogens, *S. aureus* has the highest infection incidence in the preschool age group, making it one of the earliest organisms in CF (4). Furthermore, patients infected with methicillin-resistant *S. aureus* (MRSA) experience faster CF disease progression (6-8). Currently there is no vaccine against *S. aureus*, and eradication of incident infections (especially MRSA) remains challenging (9). Hence, there is a critical need to understand how *S. aureus* initially infects the CF airway.

There is some prior evidence the *S. aureus* associates with mucus. Mucins from saliva, nasopharynx, and conjunctiva are known to bind *S. aureus* (10). In animals infected intranasally with *S. aureus*, bacteria bind to mucus from anterior and posterior turbinates without direct binding to the underlying epithelium (11). However, the molecular basis for *S. aureus* attachment to mucin is not fully understood. *S. aureus* expresses a variety of surface proteins, collectively known as adhesins, that are implicated in the pathogenesis of other serious infections like endocarditis and sepsis. We hypothesized that bacterial genetic factors regulating adhesins control *S. aureus* attachment to mucus. In this study, we observed whether *S. aureus* attaches to mucus in pig tracheas and we developed a method to quantify the affinity of *S. aureus* for mucin by a competition assay. We used this approach to measure the interaction of mucin with *S. aureus* strains with suspected defects in attachment. We examined two candidate regulators of bacterial attachment: Sortase A (SrtA), a transpeptidase enzyme required for surface display of proteins (12), and multiple gene regulator A (MgrA), a transcriptional regulator of surface adhesins, including the giant staphylococcal surface proteins Ebh and SraP (13-15).

## Materials and methods

### Bacteria strains and growth conditions

Bacterial strains and plasmids used are listed in the Table 1. *S. aureus* was grown overnight in tryptic soy broth (TSB), supplemented with 10 µg/ml of chloramphenicol (Cam) to maintain the sGFP-expressing plasmid pCM29. For construction of *S. aureus* mutants, tryptic soy agar with Cam or with tetracycline (Tet, 1 µg/ml) were also used. To construct mutant strains, bacteriophage transduction between *S. aureus* strains was performed with phage 80α or 11 as described previously (16). To generate the sortase mutant AH4480, the Δ*srtA* mutation containing a Tet^R^ cassette was transduced from the previously described Δ*srtA* mutant in the *S. aureus* strain RN4220.

**Table 1.** Bacterial Strains and plasmids used in this study.

| **Name** | **Genotype/Properties** | **Reference** |
| --- | --- | --- |
| **Strain** |  |  |
| LAC | Wild type, USA 300 CA MRSA, Erm<sup>S</sup> | Crosby (13) |
| AH3455 | LAC $\Delta mgrA$ | Crosby (13) |
| AH4480 | LAC $\Delta srtA$ | This work |
| AH3485 | LAC $\Delta mgrA \emptyset 11::LL29erm mgrA$ | Crosby (13) |
| AH4393 | LAC $\Delta clfA \Delta fnbAB$ | Deng (41) |
| AH4413 | LAC $\Delta clfA \Delta fnbAB clfB::Tn$ | Deng (41) |
| AH3798 | LAC $\Delta mrgA \Delta ebh sraP::Tn$ | Crosby (13) |
| AH3481 | LAC $\Delta mgrA \Delta ebh$ | Crosby (13) |
| AH3811 | LAC $\Delta mgrA sraP::Tn$ | Crosby (13) |
| **Plasmid** |  |  |
| pCM29 | sGFP expression vector, Cam<sup>R</sup> | Pang (42) |

### Trachea preparations

Animal studies were approved by the University of Iowa Animal Care and Use Committee. Male and female newborn pigs were obtained from Exemplar Genetics. The pigs were sedated with ketamine and acepromazine or xylazine and euthanized with I.V. Euthasol. We removed 1 cm tracheal segments following euthanasia, made a longitudinal cut on the ventral surface, and pinned the trachea flat to expose the mucosal surface. To mimic the CF electrolyte transport defect, we used conditions that impair chloride and bicarbonate transport. We submerged the tissue with 40 ml HEPES buffered saline with 10 µM bumetanide as previously described (3). 100 µM methacholine was added to stimulate submucosal gland secretion (1, 3).

### Imaging interaction of S. aureus and mucus in pig tracheas

*S. aureus* strains were grown overnight in tryptic soy broth (TSB) with 10µg/ml Cam to maintain selection of GFP plasmids (Table 1). We added 40 µL of stationary phase GFP-*S. aureus* to the 40 ml HEPES buffered saline. In some experiments, we added 1:10,000 red fluorescent nanospheres (FluoSpheres, Molecular Probes) to identify mucus strands (1, 3). We used a 25x (Nikon Apo LWD, Water, NA 1.1) objective and a Nikon A1R confocal microscope. We quantitated the GFP signal on the epithelial surface and compared to the GFP signal on mucus strands. We determined the speed of individual *S. aureus* cells on these pig tracheas using Imaris software (Bitplane) and compared the speed of free bacteria to attached bacteria.

### Quantifying the interaction of S. aureus and mucus by competition assay

Pig Gastric Mucin (type III, sialic acid 0.5-1.5%, Sigma #M1778, Lot SLBL7748V) was dissolved in phosphate buffered saline, pH 7. Mucin concentrations ranging from 4 µg/mL to 8 mg/mL were produced by serial dilution. 100 µL of diluted mucin was added to non-treated 96-well black optical plates (Thermo Fisher #265301). Methylcellulose (MP Biomedicals #155492, Lot 6257E), heparin sulfate (Sigma #H3149-250KU, Lot SLBS9674), and bovine submaxillary mucin (Sigma #M3895-1G, Lot SLBS0651V) were obtained commercially and prepared similarly as antagonists of bacterial attachment. To dissolve methylcellulose, we autoclaved it as a dry powder and dissolved it in PBS by stirring overnight at 4 °C.

For in vitro binding assays, we diluted stationary-phase GFP *S. aureus* to OD of 0.6 in PBS. Each strain of *S. aureus* had similar GFP signal per cell. 50 µL of *S. aureus* were added per well following dilution. Bacteria were allowed to attach for 1 hour at 37 °C. The mucin suspension and any bacteria not attached to the plastic surface were removed by aspiration. We performed two rounds of washing with PBS followed by suctioning. The residual fluorescence was assayed using a SpectraMaxi3X plate reader with settings optimized for GFP.

### Statistical analysis

For each experiment, we used 4 replicate wells for each condition. Following experiments, we calculated the average GFP fluorescence from replicate wells and plotted the average fluorescence versus log_10_ of the final concentration of binding antagonist measured in mg/mL. We used GraphPad Prism version 7 to perform 4-parameter logistic regression, allowing for calculation of IC_50_. We rejected experiments from further analysis if there were not at least 2 top and bottom values for fitting the logistic regression. For summary statistics, we used IC_50_ values from independent experiments performed on different days. For comparison of IC_50_ values, we used log-transformed data for statistical tests. For non-normally distributed data sets, we used non-parametric statistical tests as appropriate.

## Results

### S. aureus attaches to mucus strands in pig tracheas

In a previous study, we observed that mucus strands fail to detach after they emerge from submucosal glands in CF pig tracheas (3). If bacteria attach to these mucus strands, they may be retained in the airway. We obtained tracheas from newborn wild type pigs, submerged them in bicarbonate-free HEPES buffered saline and treated with bumetanide. These conditions block chloride and bicarbonate secretion, simulating CF airway conditions (3). We added methacholine to stimulate mucus secretion by submucosal glands. To these preparations, we added stationary phase *S. aureus* expressing GFP to test whether these bacteria associate with mucus strands, or if they bind directly to the epithelium (Figure 1). We selected LAC, a community-acquired strain of MRSA (CA-MRSA), as our model organism. CA-MRSA has recently increased in prevalence within the CF population (17).

**Figure 1.**
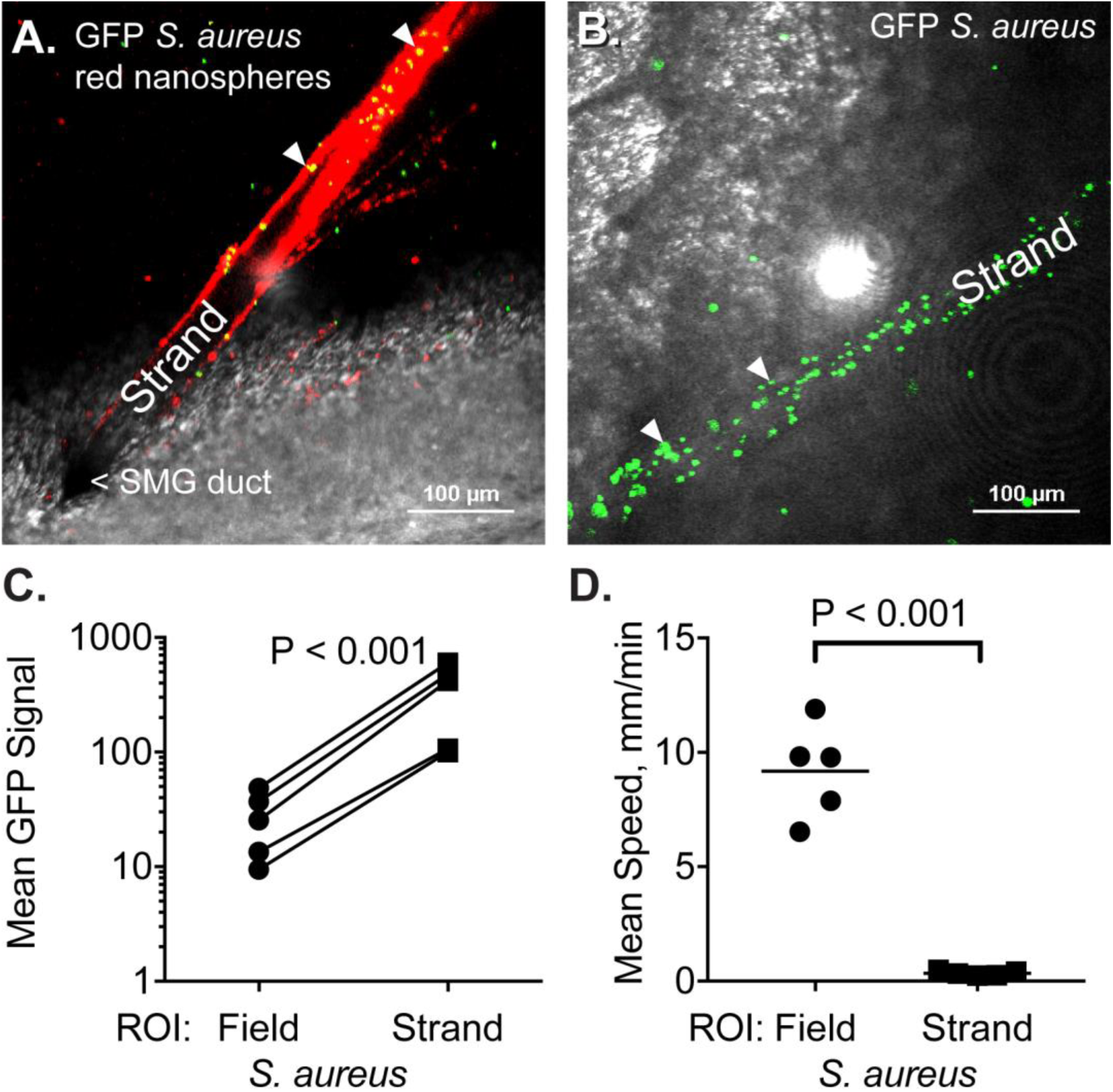
Attachment of GFP-*S. aureus* to mucus strands in pig tracheas. **A.** Confocal micrograph shows a strand of mucus that is labeled with red fluorescent nanospheres. The strand arises from a submucosal gland (SMG) duct. GFP-labeled *S. aureus* cells colocalize with nanospheres and appear yellow. Arrowheads indicate two of the attached *S. aureus* cells. **B.** Attachment of *S. aureus* to a mucus strand in the absence of nanospheres. **C.** Mean GFP signal by region of interest (ROI), either within the mucus strand or elsewhere in the field. Lines connect measurements made within the same experiment for 5 independent experiments. **D.** Mean speed of individual *S. aureus* depending on whether bacteria are attached to the strand or are elsewhere in the field. Each dot represents a unique experiment.

To help identify mucus strands, we added red fluorescent nanospheres as previously described (1, 3). In the presence or absence of nanospheres, GFP *S. aureus* was predominantly located on mucus strands (Figure 1A-C). Under these experimental conditions, mucus strands move slowly and do not properly release from submucosal gland ducts (3). *S. aureus* attached to these mucus strands had average speed less than 1 mm/min, whereas *S. aureus* that were not attached to strands moved more quickly, indicating that few were attached to the surface epithelium (Figure 1D, Supplemental Videos 1 and 2).

### Quantifying bacteria-mucus interaction by competition assay

Mucus strands form a discontinuous mesh rather than a complete gel over the airway surface (1, 2). Such a discontinuous layer may not prevent bacteria from reaching the epithelial surface. However, our confocal microscopy observations indicate that when *S. aureus* are added to pig tracheas, they accumulate on these mucus strands rather than on the epithelial surface. This might indicate that mucus strands function as competitors with the epithelial surface for bacterial binding, similar to a decoy receptor. Once bacteria attach to mucus strands, ciliary beating transports the aggregate of mucus and bacteria from the lungs. To quantitate the adherent function of mucus, we designed a competition assay to measure how efficiently standardized mucins prevent the attachment of bacteria to a standardized surface without the confounding effect of ciliary flow.

We used microtiter plates as a standardized surface for attachment. GFP-labeled *S. aureus* were added to these plates in the presence of variable concentrations of pig gastric mucin. We incubated the suspension of bacteria and mucin for one hour at 37 °C, allowing bacteria to associate with either the mucin or the plastic surface. Bacteria that did not attach to the plastic surface, but instead associated with mucin in solution, were removed by aspiration and washing. We quantitated residual bacteria attached to the surface with a fluorescent plate reader (Figure 2).

**Figure 2.**
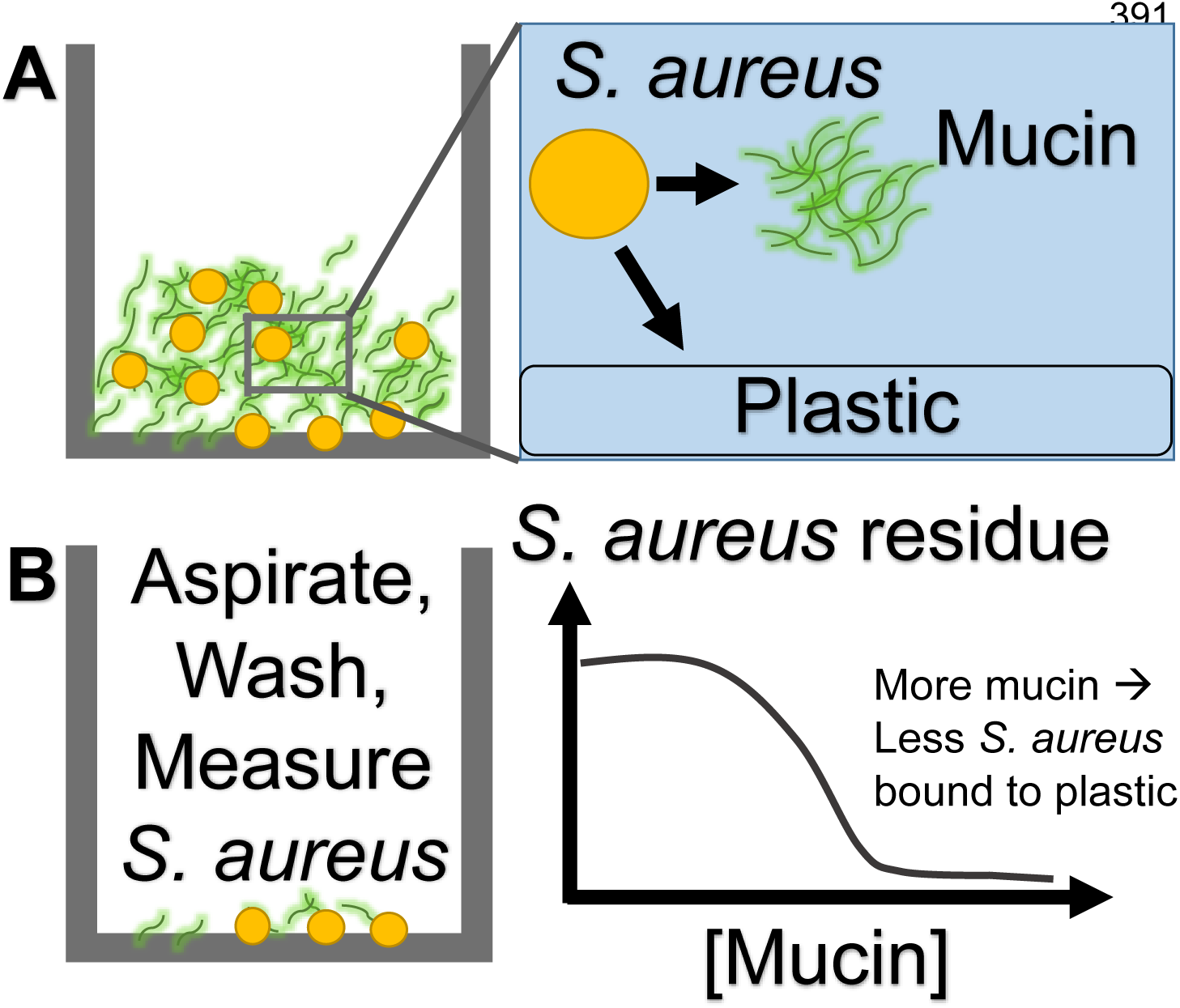
Schematic depiction of adherence assay. *S. aureus* is added to microtiter plate wells containing varying concentrations of mucin and allowed to incubate for one hour. In this scenario, the bacteria may either interact with the plastic surface or with the mucin, which functions as a competitor. After an hour incubation, the bacteria-mucin suspensions are aspirated, the microtiter plates are washed, and the remaining bacteria that are attached to the plastic are measured by their GFP fluorescence.

We first compared the attachment of wild-type LAC and mutant strains to plastic in the absence of mucin. We compared LAC to an isogenic mutant lacking Sortase A (Δ*srtA*) (12), which catalyzes attachment of *S. aureus* surface proteins to the cell wall, and MgrA (Δ*mgrA*), a transcriptional regulator of adhesin expression (13). All three strains had similar attachment to plastic prior to removal of unattached bacteria. Δ*srtA* had significantly decreased GFP signal following washing and aspiration, indicating defective attachment to plastic (Figure 3A). We did not study it further. By contrast, the GFP signal from Δ*mgrA* was indistinguishable from LAC.

**Figure 3.**
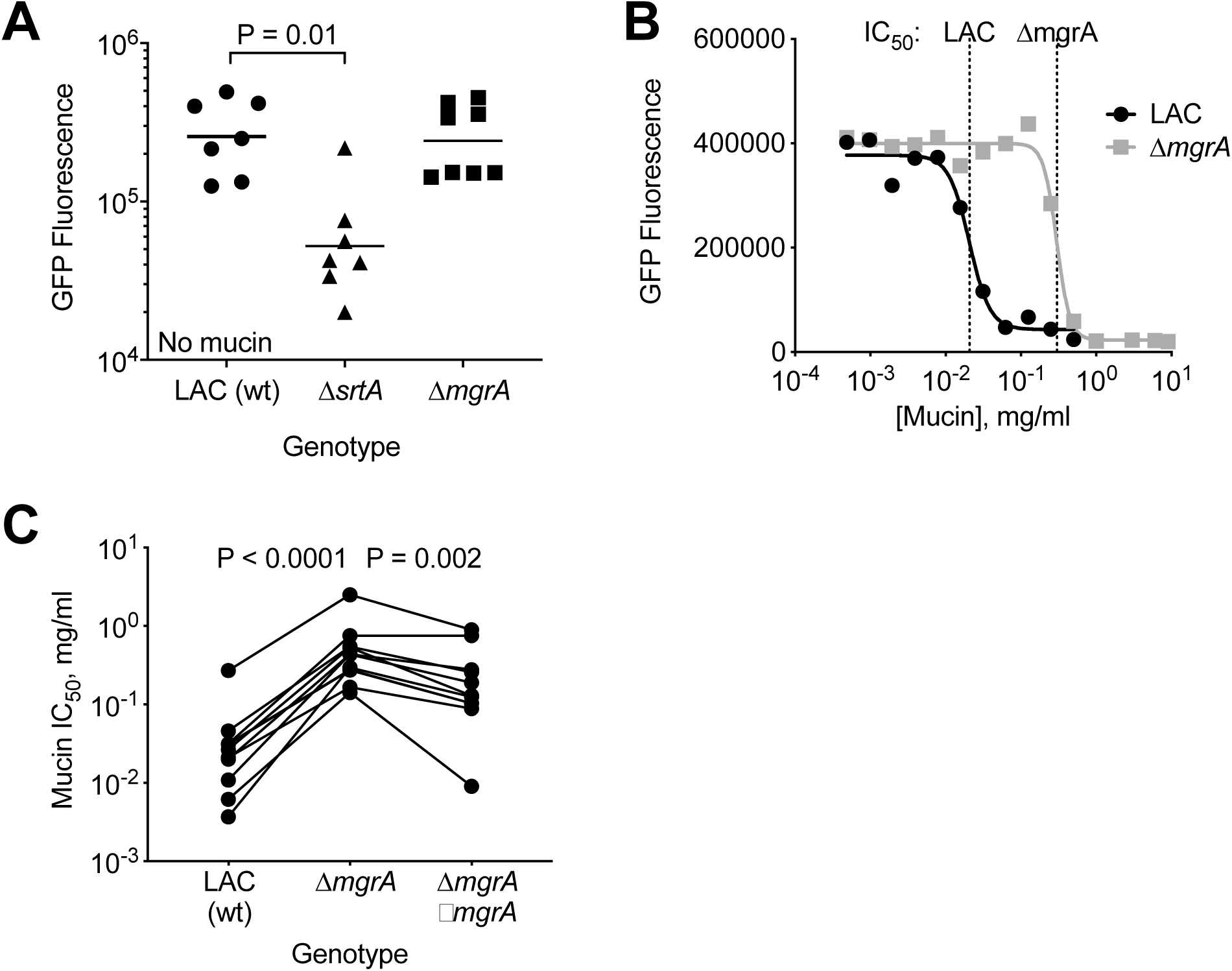
Detection of different attachment phenotypes by mucin competition assay. **A.** Attachment of *S. aureus* strains to microtiter plates as measured by residual GFP fluorescence after washing. Δ*srtA* had decreased attachment in mucin-free conditions. By contrast, Δ*mgrA* attachment was indistinguishable from isogenic LAC (wild type). Each point represents an independent experiment. *P* values were calculated by Kruskal-Wallis test, comparing mutants with LAC control. **B**. Inhibition of *S. aureus* attachment by pig gastric mucin. Mucin was serially diluted and added to 96-well plates prior to GFP *S. aureus*. After 1 hour, unattached bacteria were removed. There were dose-dependent reductions in attachment by wild type *S. aureus* (LAC, black) and a mutant deficient in MgrA (Δ*mgrA*, gray). IC_50_ estimates for mucin versus each strain are indicated at top. Data are representative of over 10 replicate experiments. **C.** Complementation of *mgrA* partially restores mucin interaction. Data show IC_50_ of mucin versus LAC, Δ*mgrA*, and Δ*mgrA* with a wild-type copy of *mgrA* integrated elsewhere on the chromosome. The IC_50_ of mucin for attachment of Δ*mgrA* was significantly higher than both wild type and the complemented strain. Complementation did not fully restore the wild-type interaction with mucin. (1-way repeated measured ANOVA, N = 8 experiments, paired by date of experiment).

### Mucin prevents bacterial attachment to plastic in a dose-dependent manner

When pig gastric mucin was added to wells with bacteria, there was a saturating dose-dependent reduction in attachment of wild-type LAC to plastic (Figure 3B). Using 4-parameter logistic regression, we determined that the IC_50_ of pig gastric mucin as an inhibitor of wild type *S. aureus* attachment to plastic was approximately 1.6 x 10^−2^ mg/mL.

### MgrA is necessary for normal mucin interactions

Using this method, we compared LAC with Δ*mgrA*, a strain with significantly altered surface adhesin expression (13). Although both LAC and Δ*mgrA* had similar binding capacity for the microtiter dish surface in the absence of mucin (Figure 3A), we observed differences in absorption to the plastic surface in the presence of mucin (Figure 3B). Significantly higher concentrations of pig gastric mucin were required to block attachment of Δ*mgrA*, as visualized by the rightward shift in the inhibition curve. We repeated these studies using a partially complemented Δ*mgrA* strain (13) and found that partial restoration of MgrA partially normalized the interaction with mucin (Figure 3C).

### MgrA-dependent differences in interaction with mucins, but not other biopolymers

To validate the observed genotype-dependent binding differences were specific to mucin, we compared adhesion of LAC and Δ*mgrA* to mucin versus relevant biologic polymers: methylcellulose, bovine submaxillary mucin, and heparin sulfate. Methylcellulose is used as a mucin substitute in pharmacologic products such as artificial tears (18). We compared it to mucin as both share viscoelastic physical properties and repeating saccharide units. Bovine submaxillary mucin is a secreted mucin with a notably higher sialic acid content than porcine gastric mucin (up to 30% vs. 1%, respectively) (19, 20, 21). We tested whether the increase in sialic acids displayed by BSM would lead to altered attachment by Δ*mgrA*, since SraP was previously shown to interact with sialic acid (22). Heparin sulfate is a linear polysaccharide abundantly secreted at mucosal surfaces. We compared it to mucin as it shares similar glycosylated residues but carries a stronger negative charge than mucin due to its sulfated moieties.

Figure 4 summarizes our results. We found the altered binding phenotype observed in the Δ*mgrA* mutant was specific to mucins. Both porcine gastric mucin and Bovine submaxillary mucin exhibited statistically significant IC_50_ differences between LAC (WT) and Δ*mgrA* strains. Conversely, methylcellulose and heparin sulfate failed to recapitulate bacterial genotypic differences in mucin adhesion.

**Figure 4.**
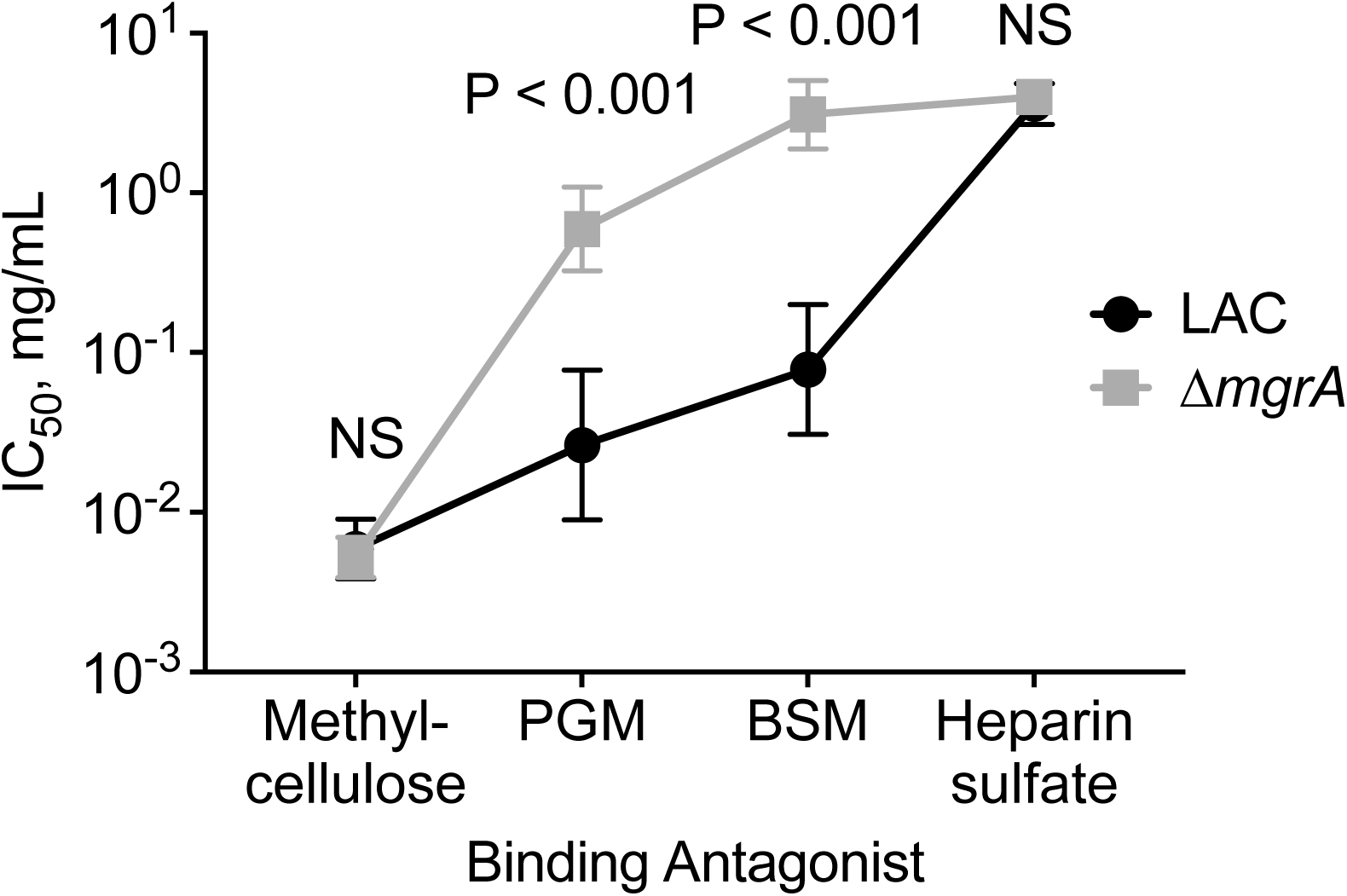
Differences between LAC and Δ*mgrA* are apparent in mucins, but not other biological polymers. Methylcellulose, pig gastric mucin, bovine submaxillary mucin, and heparin sulfate were tested for inhibitory activity against *S. aureus* attachment. IC_50_ values for each polymeric substance is shown for versus LAC (black circles) and Δ*mgrA* (gray squares). Symbols represent mean and standard error, calculated on logarithmic transformed data. Number of unique experiments per condition: 5 (Methylcellulose), 12 (PGM), 8 (BSM), and 3 (Heparin sulfate). Differences between LAC and Δ*mgrA* were observed with PGM and BSM.

These studies demonstrated that mucin is a poorer inhibitor of Δ*mgrA* attachment compared to the wild type strain LAC. We considered three possible explanations for this phenomenon. 1) MgrA is needed for the expression of classical *S. aureus* surface adhesins (13); 2) MgrA suppresses expression of surface proteins that block interaction with mucin; or 3) MgrA induces the expression of adhesins that directly bind mucin.

### A S. aureus mutant lacking conventional adhesins has normal mucin interaction

Some of the most well studied adhesins in *S. aureus* are ClfA, ClfB, and the fibronectin binding proteins A and B (FnbpAB). These adhesins have broad roles in binding fibrinogen, fibronectin, and other extracellular matrix components, and have been linked to nasal colonization, clumping and general *S. aureus* virulence properties (23). Using a mutant defective in all these adhesins (LAC Δ*clfAB* Δ*fbnAB*), we tested interactions with mucin. We found that the inhibition curve of pig gastric mucin versus this clumping factor mutant was indistinguishable from wild-type LAC (Figure 5A). Thus, the presence or absence of these important adhesins does not explain the altered interaction of Δ*mgrA* with pig gastric mucin.

**Figure 5.**
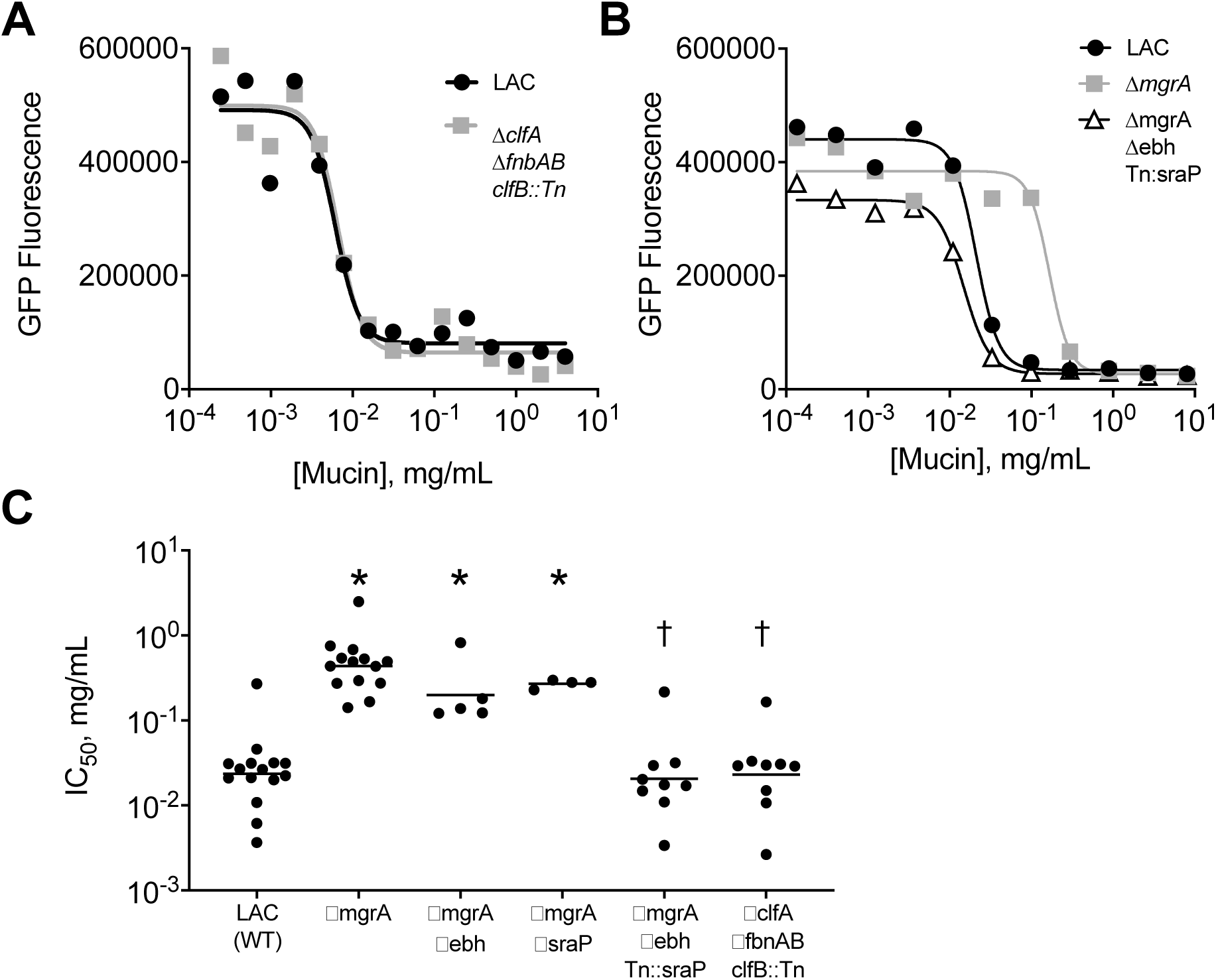
Regulation of *S. aureus* mucin interaction by adhesins downstream of MgrA. **A**. Dose-dependent inhibition of *S. aureus* attachment in the presence and absence of four classical adhesins. Representative data compare LAC (black) and an isogenic strain deficient in *clfA, clfB, fnbA*, and *fnbB* (gray). There were no statistical differences between the two strains under these conditions. **B.** Normalized interaction of Δ*mgrA* with pig gastric mucus in the absence of Ebh and SraP. In this representative experiment, pig gastric mucin blocked attachment of a triple mutant strain Δ*mgrA* Δ*ebh sraP*::Tn (white triangles) at similar concentration as was observed with the wild type LAC (black circles). Both strains had lower IC_50_ compared to Δ*mgrA* (gray squares). **C.** Summary data show IC_50_ of pig gastric mucin versus a series of *S. aureus* mutants. Each dot is an independent experiment, bars represent geometric mean. 1-way ANOVA with post-hoc comparisons to LAC or Δ*mgrA* controls. * P < 0.01 vs LAC, † P < 0.01 vs Δ*mgrA*.

### Deficiency in Ebh and SraP suppress ΔmgrA mucin interaction phenotype

MgrA functions as a transcriptional repressor that blocks the expression of the large surface proteins Ebh and SraP in the *S. aureus* LAC strain (13). We hypothesized that increased expression of one of these surface proteins was responsible for the decreased mucin binding observed in Δ*mgrA*. Therefore, we tested the interaction of mucin with Δ*mgrA* lacking either Ebh, SraP, or both (Figure 5 B and C). We found that deficiency in Ebh or SraP alone had minimal effect of mucin interaction compared to the parental Δ*mgrA* strain. However, the loss of both Ebh and SraP completely restored binding to the Δ*mgrA* strain. These data are most consistent with Ebh and SraP functioning as interchangeable factors that inhibit bacterial interaction with mucin. This is analogous to their function as anti-clumping factors (13).

## Discussion

*S. aureus* respiratory infections remain a major problem for patients with CF. It is well documented that infection with methicillin-resistant strains of *S. aureus* is associated with worsening CF disease progression (6, 7), and newer treatments for CF have had minimal effect on the prevalence of *S. aureus* (24, 25). Moreover, *S. aureus* is a major cause of respiratory disease in children and adults without CF (26, 27, 28). Although *S. aureus* may colonize asymptomatic individuals (29), *S. aureus* is a prevalent cause of community-acquired pneumonia, especially following influenza infection (30-32) and has high case fatality rates (27, 33, 34).

### *S. aureus* binds airway mucus

Because of its importance as a respiratory pathogen, it is crucial to understand how *S. aureus* initiates infections. We found that when introduced into explanted airways, *S. aureus* accumulates on strands of mucus arising from submucosal glands. When liquid secretion from these glands is blocked pharmacologically, the *S. aureus* that attach to these strands may not be cleared efficiently (3). Previous studies of *S. aureus* showed minimal contact with ciliated mucosal surface in human nasal biopsies (35) and that attachment could be decreased with lectin or monosaccharide competition (36), consistent with bacterial interaction with mucus.

Mucus is described as “sticky,” but its adhesive properties are seldom quantified. The development of a competition assay enabled us to measure the strength of the interaction between mucus and bacteria. Using the IC_50_ of competition curves as an estimate of the affinity of *S. aureus* for mucin, we found that the interaction with wild-type *S. aureus* is relatively weak (∼10^−2^ mg/mL). However, mucin production is abundant and thus it may have high capacity for removing inhaled bacteria. Since *S. aureus* can stimulate mucin synthesis (37, 38), initial infection could amplify the number of potential binding bacterial sites. This may perpetuate infection in CF, where the release of mucus is impaired (3).

### Bacterial surface factors regulate attachment and mucus interaction

How does *S. aureus* adhere to respiratory mucus? Differences between bacterial strains isolated from patients suggest genetic factors control the interaction of *S. aureus* with mucus (39). We predicted that genes regulating the display of adhesins may alter the attachment properties of *S. aureus* (23). Most of the classical surface adhesins are added to the *S. aureus* cell wall by sortase-dependent transpeptidation (12). However, we observed a deficiency in attachment to our standardized surface with Δ*srtA S. aureus* and did not study them further. MgrA regulates the expression of some adhesins on the bacterial cell wall (13, 15). In the absence of MgrA, higher concentrations of mucin were required to block attachment of *S. aureus* to microtiter plates, consistent with decreased interaction of *S. aureus* with mucin. The decreased interaction of Δ*mgrA* with mucin was not related to well-known adhesins ClfA, ClfB, or FnbpAB, as a different strain lacking these surface proteins had normal interaction with mucin. Instead, the binding phenotype of Δ*mgrA* depended on Ebh and SraP, which MgrA normally represses. We speculate that these large surface proteins sterically block the interaction of glycans from mucin side chains with low affinity receptors on the bacterial cell wall. This theory parallels recent work demonstrating decreased adhesion of Δ*mgrA S. aureus* to various host molecules, including fibrinogen, fibronectin, and collagen (15). This may also be a mechanism for the anti-clumping activity of these large bacterial surface proteins (13).

### *S. aureus* interaction with mucin may weaken with greater negative charge

Previous studies of *S. aureus* showed its attachment to mucin was only weakly inhibited by N-acetylneuraminic acid (11), and could be augmented by neuraminidase treatment (40). This implies a low affinity state between *S. aureus* and sialic acid, likely explained by charge repulsion between teichoic acids on the bacterial cell wall and sialic acid residues on mucins. We found that bovine submaxillary mucin, which has higher sialic acid content had decreased interaction with *S. aureus* compared to pig gastric mucin. Heparin sulfate, a polyanionic polymer had even weaker interaction with *S. aureus.* By contrast, there was stronger interaction of *S. aureus* with methylcellulose, a neutral polysaccharide. Future studies using *S. aureus* strains with mutations in teichoic acid biosynthesis may be useful for further testing the effect of charge repulsion between mucins and the bacterial cell wall.

## Advantages

Here, we report a quantitative, reproducible method for measuring the interaction of bacteria with mucus. This method could be adapted to studying a larger number of mutants or clinical isolates, and it could be adapted to study a variety of mucins or mucin analogs. Applying this method, we can identify *S. aureus* mutants with defective attachment to a standard surface and can quantitate the relative affinity of different *S. aureus* strains for mucin.

## Limitations

The method we report does not capture the complexity of an *in vivo* or *ex vivo* airway epithelium, in which antimicrobial proteins, innate immune cells, and an abundance of macromolecules other than mucin may influence bacterial colonization. Furthermore, we did not replicate the cilia-directed flow and continuous secretion of mucin seen in airway infections. More specifically to mucin, pig gastric and bovine submaxillary mucins are standardized, commercially available mucin glycoproteins, but they have been denatured and do not retain their original viscoelastic properties. On the other hand, airway mucins are highly ordered (1, 2) and possess their native molecular properties. Our competition method for measuring interaction of mucin and bacteria is indirect and requires intact attachment of bacteria to the microtiter plate. However, because these mucins are easily washed off the plate surface, we found the indirect method was more feasible than direct interaction studies and did not require additional steps such as chemical modification of the mucins to conjugate them to a surface.

## Conclusions

Our study shows that *S. aureus* attaches to strands of mucus. These strands are poorly cleared in CF. With a competition assay, we show that the affinity of *S. aureus* for mucin is relatively weak (10^−2^ mg/mL) and may be regulated by MgrA through repression of Ebh and SraP. This method will enable further study of bacterial factors involved the interaction with host mucins.

## Acknowledgements

This work was funded in part by NIH K08 HL136927 (AJF), CFF FISCHE16I0 (AJF), and NIH AI083211 (ARH). CPP was supported a fellowship for medical students NIH T35 HL007485 40. We thank Michael Welsh for his constructive feedback at several stages of this project.

## Author Contributions

Conceived and designed studies: AJF, ARH

Performed experiments: CPP, CRS, PDA, NA, ALC

Wrote first draft of paper: CPP, AJF

Edited paper: CPP, AJF, ARH

**Supplemental Video 1.** Confocal microscopy of excised pig trachea demonstrates reduced speed of *S. aureus* attached to mucus. Pig tracheas were treated with bumetanide while submerged in buffer lacking bicarbonate, slowing the emergence of mucus. Red fluorescent nanospheres were added to identify demonstrate mucus strands, which emerge from submucosal gland ducts. Green color indicates *S. aureus* that express green fluorescent protein. Unattached *S. aureus* are transported by ciliary flow. Many *S. aureus* are attached to the mucus strand. Their movement is slow compared to the unattached bacteria. Video is shown in real time.

**Supplemental Video 2.** Confocal microscopy of excised pig trachea shows an attachment event by *S. aureus*. Pig trachea was prepared similarly to the previous video, with red fluorescent nanospheres used to illuminate mucus strands. GFP expressing *S. aureus* were added. The video pauses at 10 s, when a new green particle of *S. aureus* appears immediately below the gland duct orifice. The video is slowed to 25% of normal speed until the *S. aureus* appears on the mucus strand (20 s). The *S. aureus* subsequently remains on the mucus strand.

